# Evaluation of window fullness at time of eclosion as an operational welfare indicator in farmed black soldier flies (*Hermetia illucens*)

**DOI:** 10.64898/2026.11.15.766814

**Authors:** S.O. Durosaro, M. Barrett

## Abstract

Body condition scoring can serve as an operational indicator of an animal’s current welfare state and may also shed light on preceding, welfare-relevant conditions, such as stress. In farmed insects, limitations posed by the exoskeleton largely preclude traditional body condition assessment. However, adult black soldier flies (BSF) farmed in the insect protein industry have transparent abdominal panels that allow for visualisation of a portion of their fat body. Prior work has shown that the relative size of the fat body in the window, termed ‘window fullness’ (WF), at time of eclosion was reduced in adult females that were handled daily as prepupae (which also led to mortality, reduced body size, and developmental delays), suggesting it may predict prior stress. Accordingly, we investigated if WF correlated with relative lipid levels in newly emerged BSF; determined if assessors could score WF on an ordinal scale by eye; and assessed if WF changed when flies were pupated in non-preferred substrates. Relative lipid content strongly correlated with WF in males but not in females. Across all observers, WF could be scored by eye on an ordinal scale with 78.38% accuracy (range: 70.27 – 89.19%). Pupating in a preferred substrate correlated with increased dry mass and head widths for both sexes; females had increased WF (14.81%) but male WF was not affected. Overall, these data suggest WF could have promise as an on-farm-assessable indicator of body condition for BSF; however, there may be sex-specific differences in validity or reliability that require further investigation.

## Introduction

Body condition score is a well-established and widely used animal-based indicator in the assessment of vertebrate farm animal welfare, having been used with pigs (Renggaman *et al*. 2015), goats (Vieira *et al*. 2015; Ghosh *et al*. 2019), cattle (Roche *et al*. 2009; Matthews *et al*. 2012; Nielsen *et al*. 2025), and sheep (Zufferey *et al*. 2021). It is commonly used because it is inexpensive, non-invasive, and can be easily performed in the field or on farms (Welfare Quality® 2009; Vall *et al*. 2025), with poor body condition suggestive of poor current welfare as well as prior, welfare-relevant conditions like starvation or stress (Welfare Quality® 2009; Veissier *et al*. 2012; Fisher *et al*. 2020; Lindholm-Perry *et al*. 2024). However, despite its widespread use, body condition scoring also has numerous weaknesses as a welfare indicator, including only serving as an indicator of poor welfare, not measuring the animal’s actual affective or emotional state, and its insensitivity to acute welfare challenges (Matthews *et al*. 2012; Fisher 2020; Fisher *et al*. 2020).

Although there are weaknesses in the use of body condition scoring to assess welfare, it remains a part of many welfare assessment frameworks as a rapid and easy to assess operational indicator for severe, chronic welfare challenges in farmed vertebrates (Welfare Quality® 2009).

However, body condition scoring is not typically employed with adult insects as their rigid and chitinous exoskeleton largely precludes the visualisation of internal body structures and, thereby, body fat levels (Johnson & Barrett 2025). However, fat levels are still expected to be relevant to insect fitness. Fat is essential for fly survival under energetic stress (Ballard *et al*. 2008, Lee & Jang 2014), providing energy during times of feed scarcity (which is common in adult black soldier flies, which are not fed on farms; Barrett *et al*. 2023; Barrett *et al*. 2025; Durosaro & Barrett 2025). Lipids also support reproductive success in *Drosophila melanogaster* (Diptera: Drosophilidae) flies, as females with more lipid reserves have been shown to lay more eggs (Kalra & Parkash 2014).

The black soldier fly (BSF, *Hermetia illucens,* Diptera: Stratiomyidae) is a farmed insect species of neotropical origin that is now reared all over the world (Marshall *et al*. 2015; Kaya *et al*. 2021; Nayak *et al*. 2024). It is farmed for waste management (Amrul *et al*. 2022), animal feed (Dabbou *et al*. 2018; Nogales-Mérida *et al*. 2019), organic fertilizer (Borkent & Hodge 2021), oil (Kim *et al*. 2020), and biodiesel production (Leong *et al*. 2016). This species is considered the ‘crown jewel’ of the booming insects as food and feed industry because of its high nutritional value, exceptional waste conversion ability, rapid growth and reproduction, and lower environmental footprint than vertebrate protein when reared on waste (Tomberlin & Van Huis 2020). It is farmed in the trillions across the globe (McKay & Shah 2025), with juveniles (larvae) slaughtered as the protein product and adults used for breeding and egg laying.

The BSF lifecycle involves multiple stages: egg, larvae (actively feeding), prepupae (non-feeding), pupae (metamorphosing), and finally adult (Tomberlin *et al*. 2002). Prepupal flies are sometimes called the ‘wandering’ stage, as they actively leave their feeding substrate and search for a safe place to pupate. BSF prepupae dig into dry substrates for pupation: this substrate will protect them from predators, parasites, and desiccation during the immobile pupal period (Rivers *et al*. 2011; Georgescu *et al*. 2020; Kortsmit *et al*. 2023). After 8-10 days as pupae, adults will then emerge (called ‘eclosing’).

Unlike many insects, body condition scoring may be possible in adult BSF. This is because adult BSF have two transparent abdominal panels through which part of their fat body can be easily visualised (Oliveira *et al*. 2016; Harjoko *et al*. 2023). Some prior research suggests that newly-eclosed adult fat body size, as visualised through this window (termed ‘window fullness’, WF), may be impacted by pre-adult stress in BSF. For instance, WF decreased on the day of eclosion in adult female flies that had been exposed to disturbance stress as prepupae; this stress also caused reduced body sizes, prolonged development times, and increased mortality (Durosaro *et al*. 2026). WF can be impacted by age in BSF (Harjoko *et al*. 2023), making it important to standardise assessment periods to early adulthood if the indicator is meant to be used to potentially assess pre-adult stress.

One form of sublethal juvenile stress may come from inappropriate pupation substrates. Various pupation substrates, such as potting soil, sand, vermiculite, manure, wood shavings, and corn cob grit, are used in BSF rearing (Holmes *et al*. 2013; Dzepe *et al*. 2020; Boafo *et al*. 2023; Liu *et al*. 2023), with no records of the substrates used by prepupae/pupae in the wild to give guidance on natural behavior. Pupation substrates seem to affect animal fitness: for instance, adult emergence rates of 92%, 69%, and 79% were reported for wood shavings, fine sand, and wheat bran, respectively, by Dzepe *et al*. (2020). Durosaro *et al*. (2026) showed further variation in survival and adult fly body size across five different substrates. However, other studies have shown substrates like sand, topsoil, and wood shavings to have high and equal pupation rates of 95-98% (Holmes *et al*. 2013) or 88-92% (Dzepe *et al*. 2020). Despite the likely importance of this substrate to this BSF life stage, no studies have provided a choice between substrates to BSF prepupae to determine if they have preferred substrates or if preference predicts fitness. When animals are provided with the opportunity to choose what they prefer, it can improve their welfare by providing them with control over their environment, ultimately reducing stress, increasing positive affective states, and increasing expression of their species-specific natural behaviors (Englund & Cronin 2023).

Accordingly, the goals of this study were two-fold. First, we hypothesised that prepupae would have substrate preferences and that these preferences would correlate with improved, fitness-relevant metrics like body size and emergence rate for adults. We predicted that preferred substrates would lead to increased adult emergence rates and larger body sizes in both males and females. Second, we hypothesised that sublethal juvenile stress from pupating in non-preferred substrates would alter WF in adult flies. We predicted that forced pupation in non-preferred substrates would reduce WF in emerging adults. We also assessed the feasibility of using WF as a measure on farms by having multiple observers with variable levels of familiarity with BSF score WF by eye and comparing agreement between observers - and between observers and more accurate, photographically-determined WF - under a microscope. Finally, we assessed if WF correlated with total body lipid content in male and female BSF.

## Materials and Methods

### Animal sourcing and husbandry

The BSF prepupae used in this experiment were obtained from Fluker Farms (Port Allen, LA, USA) as larvae and allowed to prepupate in Gainesville diet (Hogsette 1992). Only darkly sclerotised and mobile prepupae were used in all the experiments.

### First pupation substrate preference test

Three pupation substrates were used for pupation substrate preference test: sand (SD; Pavestone natural play sand), potting soil (PS; Miracle.Gro indoor potting mix), and wood shavings (WS; So phresh natural aspen small animal bedding). WS were dry when purchased; SD and PS were dried in an oven (VWR convection oven, Germany) at 100 °C for 24 hours.

A plastic card was used to divide 32 oz containers (height: 14 cm, lower diameter: 8.5 cm, and upper diameter: 12 cm) into two equal sides. Each side was loosely filled with a pair of different substrates (PS v. SD, PS v. WS, and SD v. WS) to a height of 10 cm. 10 mL of water was added to each substrate, and the divider was gently removed. Fifteen prepupae were placed in the middle of the containers on the dividing line between the two substrates. A single paper towel was placed on the top of the containers and tied around the container with flagging tape to prevent the prepupae from escaping but allow for ventilation. Each substrate pair condition was replicated six times (n = 6 cups/pairing; n = 270 total prepupae). All cups were placed in an incubator (Percival I-36NL, USA) with a bin of water at the bottom to increase RH. Temperature was 24.49 ± 0.01 °C and RH was 87.07 ± 0.13%; a 14:10 L:D cycle was supplied by an 8.5 Watt (800 Lumen) overhead LED bulb. Replicates of a substrate pairing were assigned so that 50% had substrate one on the left, and 50% had substrate one on the right, to account for any side bias.

On day 8, the substrates on each side of the cup were slowly removed with a laboratory spatula, and the substrates in which each prepupa or pupa was found were recorded. The depth at which each mobile or non-mobile prepupa or pupa was found was measured from the original substrate height. Only non-mobile prepupae and pupae were used in analysing the pupation preference index and depth. The pupation substrate preference index was calculated as:

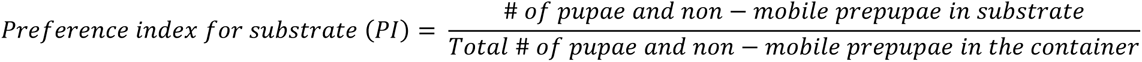

### Effect of exposure to light on preference for WS

In the first preference test, WS were strongly non-preferred; we thought this could be due to the looser texture of WS letting in more light as BSF larvae and prepupae are expected to be photophobic. Therefore, to determine if light exposure caused the reduced preference for WS, we performed another experiment similar to the first preference test, but only WS and SD were used as pupation substrates. Twelve containers that were WS v. SD were created. The exterior lateral surfaces and the bottom of half of the containers were covered with black, opaque aluminium Cinefoil. The six containers covered with black, opaque aluminium foil were also placed underneath a box, and all the containers were placed in the incubator with the conditions and data collection otherwise the same as the first preference test.

### Effect of preferred and non-preferred substrate on life history traits

To determine if substrate preference affects life history traits such as eclosion (time and rate), mass (adult wet and dry mass), morphology (head width and thorax length), and WF in BSF, another experiment was set up with prepupae allowed to pupate in the preferred and non-preferred substrates. Briefly, 32 oz plastic containers were filled with each of the preferred (PS) and non-preferred substrates (WS) to a depth of 10 cm. 20 ml of water was added to each substrate. Fifteen darkly sclerotised mobile prepupae were placed on top of the substrates, and a single paper towel was placed on the top of the containers and tied around the container with flagging tape to prevent the prepupae from escaping before pupation. The preferred and non-substrates were replicated four times each (n = 120 total prepupae). All the 8 containers were placed in an incubator (Percival I-36NL, USA) set at room temperature/RH (not measured, but see first pupation substrate test for general climate levels in the lab) and 14:10 L:D cycle supplied by an 8.5 Watt (800 Lumen) LED bulb.

Life history traits were studied as in Durosaro *et al*. (2026). Briefly, each container was checked for adult emergence once every day until day 26. When adults were found, eclosion date was recorded. Then, adults were removed from the containers, anesthetised with isoflurane, sexed, and weighed (for wet mass) to the nearest 0.1 mg with an analytical scale (VWR-220TC). The ventral abdominal window of each fly was photographed against a red background under a microscope (AMScope, SM-1TSZ-V03) at 2X magnification with a 0.5X camera mount. The thorax length of the adult flies was measured as the distance between the midpoints of prescutum and the end of scutellum to the nearest 0.1 mm using a digital caliper (WEN). The head width was measured as the distance between the widest parts of the head using a digital caliper (WEN) to the nearest 0.1 mm. After measuring the thorax length and head width of the flies, they were frozen for at least 24 h at -20 C for euthanasia while still anesthetised. Then, flies were placed individually in 1.5 mL Eppendorf tubes with the cap opened in an oven (VWR convection oven, Germany) at 60 °C for 72 hr, and the dry mass was recorded to the nearest 0.1 mg after removal from the oven.

The photos of the fly’s ventral abdominal windows were analysed using ImageJ (Schneider *et al*. 2012) to quantify the fat body and total window areas. WF was calculated as fat body area divided by total body area (Harjoko *et al*. 2023). Eclosion rate was calculated as the number of adults that emerged from a container divided by 15 (the number of prepupae put in a container) multiplied by 100.

### Inter-observer reliability of by-eye WF assessment

The fat content of the abdominal window appears as an opaque, creamy bluish to whitish mass (Figure 1A-E; photos taken from a prior experiment, Durosaro *et al*. 2026). To determine if WF can be reliably scored without the use of a microscope, four observers with different levels of BSF experience were briefly trained (shown in Supplementary File 1) to visually score BSF WF according to five scores using photographs of WF from another experiment:

- score 0: window 0-19.99% full of fat body (Figure 1A)
- score 1: 20-39.99% full of fat body (Figure 1B)
- score 2: 40-59.99% full of fat body (Figure 1C)
- score 3: 60-79.99% full of fat body (Figure 1D)
- score 4: 80-100% full of fat body (Figure 1E).

**Figure 1A-E.**
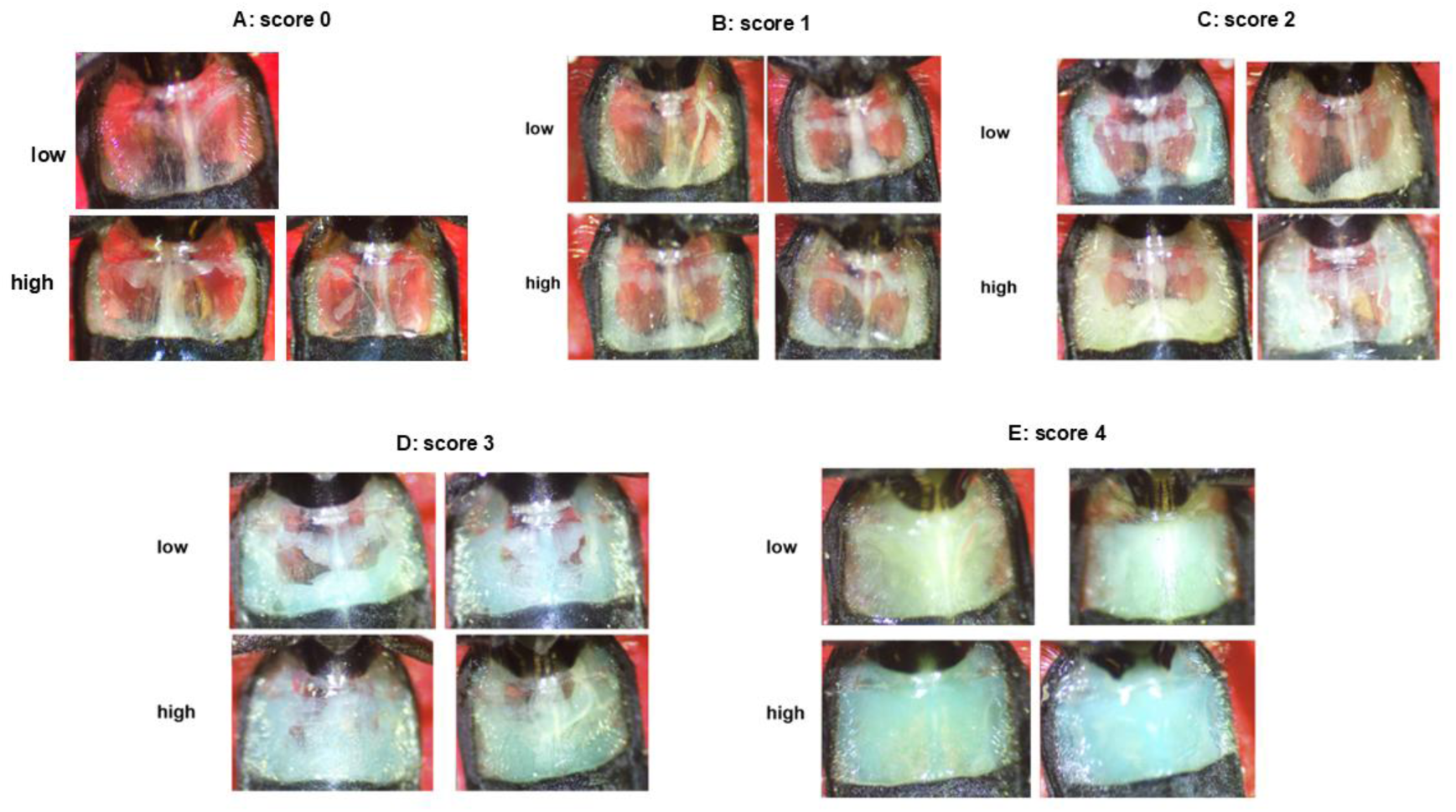
WF according to 5 ordinal scores. (A) score 0: window 0-19.99% full of fat body (B) score 1: 20-39.99% full of fat body (C) score 2: 40-59.99% full of fat body (D) score 3: 60-79.99% full of fat body (E) score 4: 80-100% full of fat body

Observer 1 had previously photographed and used ImageJ in scoring the WF of more than 1000 BSF and had five years of BSF experience. Observers 2 and 3 have previously worked with BSF but never photographed or used ImageJ in scoring the WF of BSF before, and had 1 – 2 years of BSF experience. Observer 4 had seen photos of BSF windows before when assisting in the laboratory, but had never scored windows and had only hours of BSF experience. BSF WF was scored live, but anesthetised with isoflurane, by each observer.

Observers were then tasked with independently scoring the WF of 37 flies that emerged from preferred and non-preferred pupation substrates using only an illuminated loupe (Ledlight, 10X – 21mm). The WF of each of these 37 flies was also photographed and quantified, blind to individuals, using ImageJ.

### Correlation between WF and total body lipid content

To determine if WF is a true measure of BSF lipid content, the lipid content and the WF of another newly emerged 48 flies from another experiment were also determined. The WF was determined as described earlier. The lipid content was determined using the method described by Williams *et al*. (2011). Briefly, flies that eclosed within 24 hours were collected, anesthetised with isoflurane, weighed, and their window photos taken. The flies were then placed individually in 1.5 mL Eppendorf tubes with the cap closed and submerged in liquid nitrogen and stored at - 80 °C before lyophilisation. The flies were later lyophilized using a freeze dryer (Harvest Right Scientific Benchtop v5. 5S). After lyophilisation, the flies were weighed on an analytical scale to obtain their dry mass. After obtaining dry mass, the flies were placed individually in 1.5 mL Eppendorf tubes, 1 mL of diethyl ether was added to each tube, removed, and replaced every 24 hr for 3 days. On the 4^th^ day, the diethyl ether was removed, and the flies were placed in a convection oven at 60 °C for 24 hr. The lipid content was calculated as the difference between the dry mass after lyophilisation and the dry mass after removing the flies from the oven.

### Statistical analysis

All analyses were performed in R (version 4.4.2) and visualisation was performed in R (version 4.4.2) or GraphPad Prism (version 10.6.0). Data normality was checked using Shapiro-Wilk test; alpha was set at 0.05 for all analyses.

The difference in pupation substrate preference index for every substrate pair was analysed using a chi-square test against the 50:50, no-preference null hypothesis. To determine if darkness changed preference for WS, the PI for WS in light was compared to the PI for WS in dark using a chi-square test against the 50:50, no-preference null hypothesis.

Data were analysed for males and females separately in the no-choice life history trait tests because BSF show significant sexual dimorphism. The difference between eclosion time, wet mass, dry mass, head width, and thorax length of flies that pupated in the no-choice preferred and non-preferred substrates was analysed with a generalised linear mixed model using substrate as the main factor, while container and average prepupal mass of each replicate were used as random factors. Poisson distribution and log link function were used in the analysis of eclosion time because eclosion time was recorded as a whole day and was not normally distributed. Gamma distribution and log link function were used in the analysis of wet mass and dry mass because both were positive, continuous, and not normally distributed. Gaussian distribution and identity link were used in the analysis of head width and thorax length because they were continuous and normally distributed.

The difference in eclosion rate and the WF of flies that pupated in preferred and non-preferred substrate was analysed using beta regression, using substrate as the main factor, while container and average prepupal mass of each replicate were used as random factors.

The inter-observer reliability across all four observers was determined using intraclass correlation coefficient [ICC (3, k)]. The agreement patterns were visualised using frequency heatmaps of true scores against the observer’s scores. The agreement between each of the four observers and the true WF score was determined using intraclass correlation coefficient [ICC (3, 1)] and weighted Cohen’s kappa. The accuracy of each observer was determined using mean bias, root mean square error, and Spearman correlation. All analyses were performed in R (version 4.4.2).

The correlation between determined lipid content and WF was analysed with Spearman correlation, and a linear model was fitted to generate a regression equation between the two variables.

### Ethics Statement

Experiments on insects are not required to be approved by the Institutional Animal Care and Use Committee in the United States of America, therefore no ethical approval could be obtained for this study. However, we applied the 3R (replacement, reduction, and refinement) principles as much as possible. We calculated appropriate sample sizes for our effect size of interest *a priori* using G*Power (power = 0.8, alpha error probability = 0.05). For refinement, we anesthetised flies with isoflurane before handling and euthanasia, which was conducted by freezing flies at (−20 °C). Flies were ordered in from a producer as late-stage larvae in order to eliminate the need to rear the full colony cycle, this minimising the overall number of animals used.

## Results

### Pupation substrate preference

BSF prepupae preferred PS and SD alike for pupation (χ^2^ = 0.76, p = 0.38, Figure 2A). However, the prepupae preferred PS and SD to WS for pupation (both χ^2^ > 65.64, p < 0.0001). The mean preference index difference between PS and WS was 0.86 (Figure 2B) and between SD and WS was 0.95 (Figure 2C).

**Figure 2A-C:.**
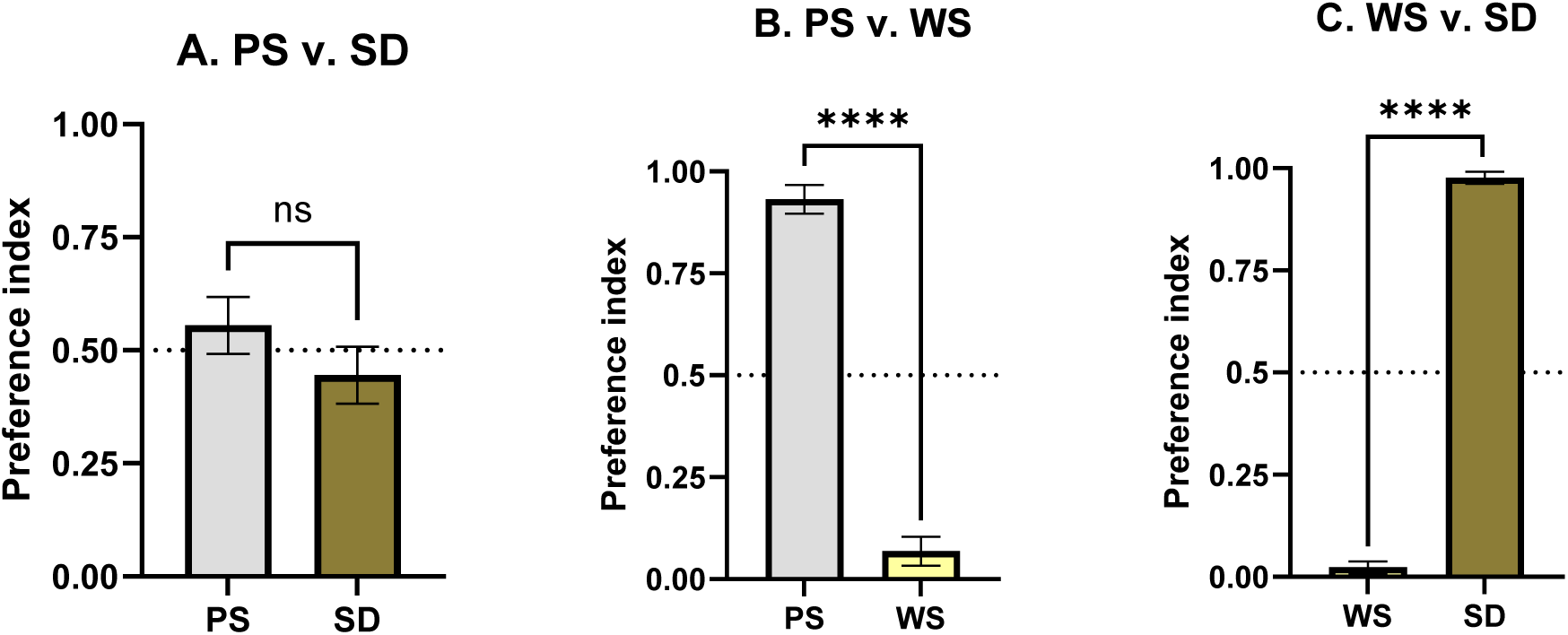
Pupation substrate preference index when BSF prepupae were put in a container containing (A) PS and SD (B) PS and WS (C) WS and SD. The BSF prepupae preferred PS to WS, SD to WS, but preferred PS and SD alike. Mean ± SE of preference index reported. ns = not significant, **** = p < 0.0001

When given the choice between WS and SD in the second experiment, there was no difference in the preference index for WS when the BSF prepupae were kept in total darkness and non-dark conditions (χ^2^ = 0.32, p = 0.57, Figure 3A). BSF prepupae preferred SD to WS under total dark (χ^2^ = 42.88, p < 0.0001, Figure 3B) and non-dark (χ^2^ = 55.85, p < 0.0001) conditions.

**Figure 3A-C:**
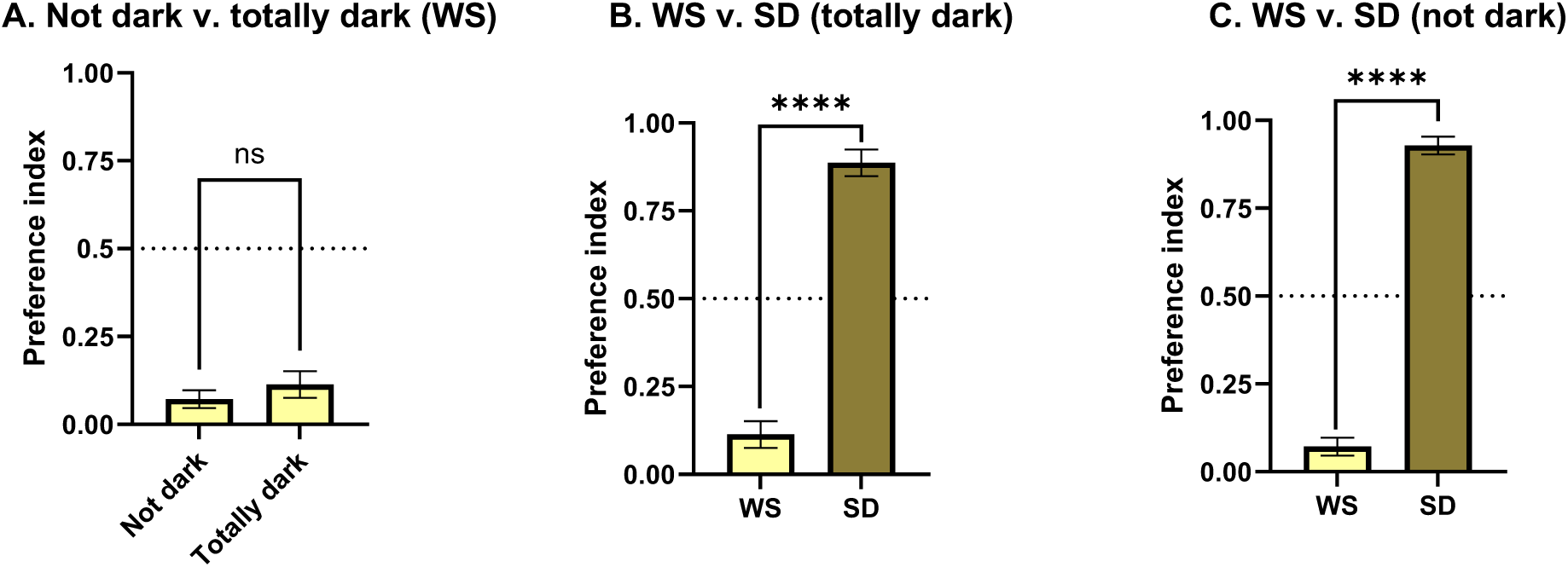
Pupation preference index (A) for WS when BSF prepupae were kept under non-darkness and total darkness (B) for WS and SD when kept in total darkness (C) for WS and SD when kept in non-darkness. There was no difference in the preference index for WS under light and in total darkness. BSF prepupae preferred SD to WS under total dark and non-dark conditions. Mean ± SE of preference index reported. ns = not significant. **** = p < 0.0001.

The number of prepupae that pupated in WS was very small (n = 6 for PS v. WS and n = 2 for WS v. SD). We therefore only analysed the pupation depth data using descriptive statistics. We found: (1) a mean pupation depth of 2.17 ± 0.27 cm for PS and 3.66 ± 0.36 cm for SD (in the PS v. SD choice test); (2) a mean pupation depth of 3.54 ± 0.28 cm for PS and 4.95 ± 1.22 cm for WS in PS v. WS choice test; and (3) pupation depth of 1.40 ± 0.99 cm for WS and 3.67 ± 0.24 cm for SD in WS v SD choice test. No clear patterns emerged in substrate pupation depth based on substrate type.

### Effect of pupation substrate on eclosion, mass, WF, and morphology

Pupating in a preferred v. not preferred substrate (PS v. WS) had no effect on eclosion rate (χ^2^ = 0.98, p = 0.32) and eclosion time (males: χ^2^ = 0.32, p = 0.57, females: χ^2^ = 0.20, p = 0.66) of flies (Table 1).

**Table 1.**
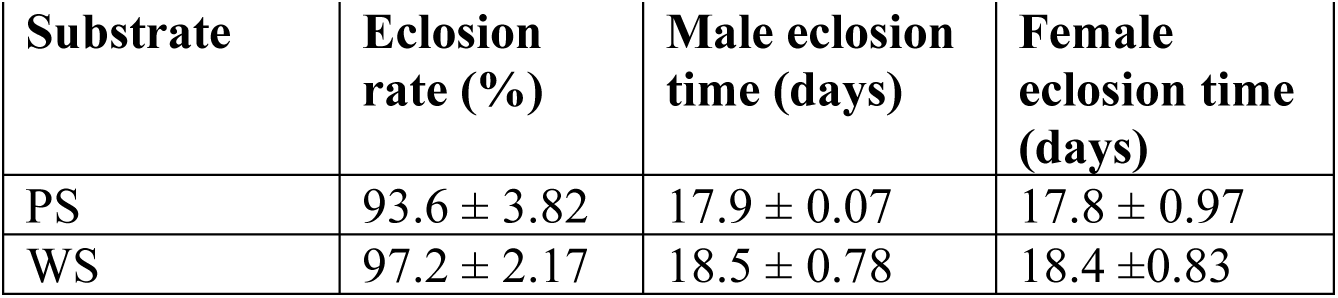
Eclosion rate and time of flies that pupated in preferred (PS) and non-preferred (WS) substrates.

Pupation substrate caused a marginal reduction in the wet mass of male flies (χ^2^ = 3.64, p = 0.06) and had no significant effect on wet mass of female flies (χ^2^ = 2.28, p = 0.13, Table 2). However, WS reduced dry mass of males by 15% (χ^2^ = 10.42, p = 0.001) and females by 18.2% (χ^2^ = 9.47, p = 0.02) compared to flies that pupated in PS. Pupation substrate did not affect WF in male flies (χ^2^ = 0.11, p = 0.74) but WS reduced WF in female flies by 14.81% (χ^2^ = 4.75, p = 0.029; Table 2, Figure 4).

**Figure 4:**
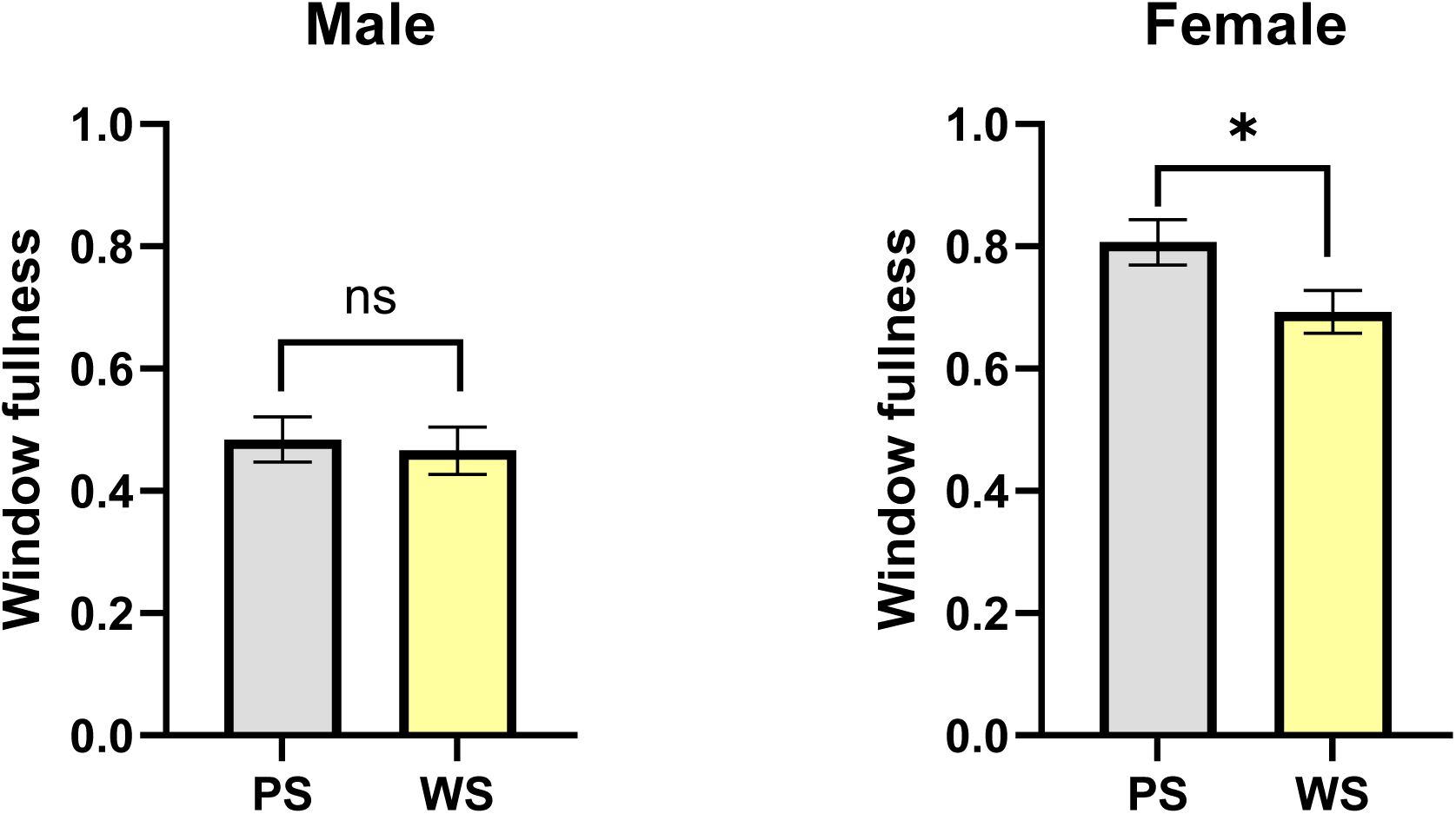
Window fullness of flies that pupated in preferred (PS) and non-preferred (WS) substrates. Pupation substrate did not affect WF in male flies, but WS reduced WF in female flies by 14.81%. Mean ± SE of preference index reported. * = P < 0.05, ns = not significant.

**Table 2.** Wet and dry mass, as well as WF, of flies that pupated in preferred (PS) and non-preferred (WS) substrates.

| **Substrate** | **Male wet mass (g)** | **Female wet mass (g)** | **Male dry mass (g)** | **Female dry mass (g)** | **Male WF** | **Female WF** |
|---|---|---|---|---|---|---|
| PS | 0.056 ± 0.002 | 0.057 ± 0.003 | 0.020 ± 0.001<sup>a</sup> | 0.022 ± 0.001<sup>a</sup> | 0.48 ± 0.04 | 0.81 ± 0.04<sup>a</sup> |
| WS | 0.051 ± 0.002 | 0.052 ± 0.002 | 0.017 ± 0.001<sup>b</sup> | 0.018 ± 0.001<sup>b</sup> | 0.47 ± 0.04 | 0.69 ± 0.04<sup>b</sup> |

<sup>a,b</sup>Letters indicate statistically significant differences between treatments within same column.

WS also reduced the head width of males (χ^2^ = 10.72, p = 0.001) and females (χ^2^ = 4.48, p = 0.034) by 3.42% and 3.36%, respectively (Table 3). There was no difference in the thorax length of flies that pupated in PS v. WS (males: χ^2^ = 0.02, p = 0.88, females: χ^2^ = 1.56, p = 0.28, Table 3).

**Table 3.** Morphology of flies that pupated in preferred (PS) and non-preferred (WS) substrates.

| **Substrate** | **Male head width (mm)** | **Female head width (mm)** | **Male thorax length (mm)** | **Female thorax length (mm)** |
|---|---|---|---|---|
| PS | 3.51 ± 0.03<sup>a</sup> | 3.57 ± 0.04<sup>a</sup> | 4.79 ± 0.07 | 4.67 ± 0.08 |
| WS | 3.39 ± 0.03<sup>b</sup> | 3.45 ± 0.04<sup>b</sup> | 4.80 ± 0.07 | 4.55 ± 0.07 |

<sup>a,b</sup>Letters indicate statistically significant differences between treatments within same column.

### Observer agreement and accuracy of scoring WF

A 60.14% accuracy was observed across all observers when scoring the WF. Overall, underscoring (30.41%) was more common than overscoring (9.46%). The majority of mismatches were off by only one score level. All observers scored 2 and 3 interchangeably (Figure 5A), indicating that the observers had difficulty differentiating between scores 2 and 3, so scores 2 and 3 were collapsed to a single score.

**Figure 5A:**
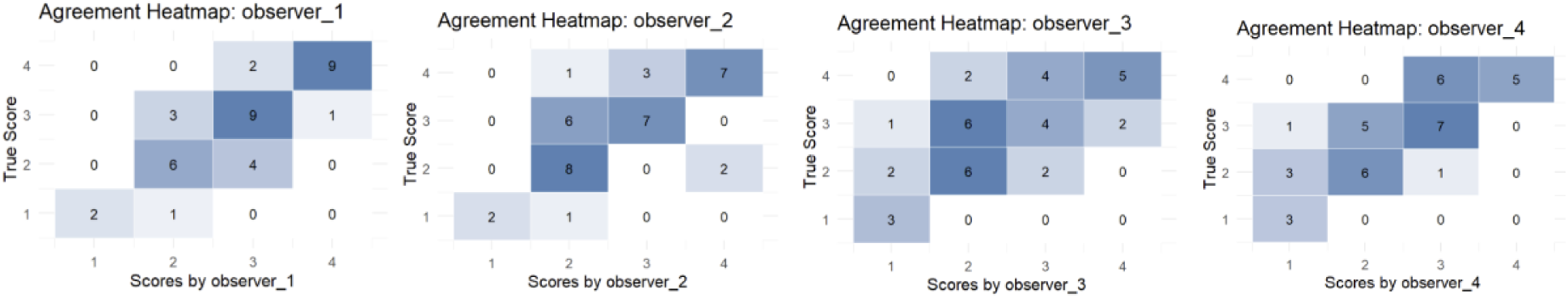
Agreement heatmaps comparing true WF scores with observers’ scores when scores 2 and 3 were not collapsed. The diagonal represents perfect agreement between the true score and observers’ scores. Observers were not able to easily differentiate between scores 2 and 3.

After collapsing scores 2 and 3 into a single score, observer agreement with the true score improved substantially. Across all the four observers, the exact agreement with true scores increased to 78.38%, underscoring reduced to 16.89% while overscoring reduced to 4.73% (Figure 5B).

**Figure 5B:**
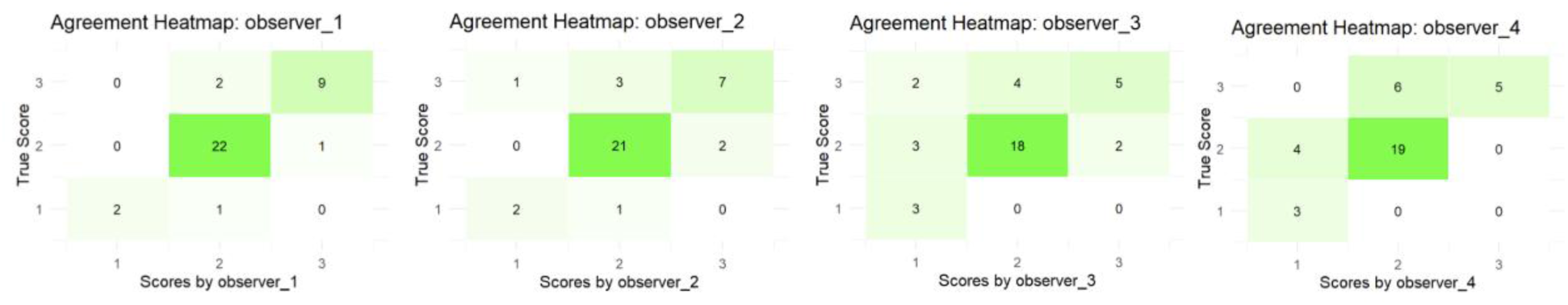
Agreement heatmaps comparing true WF scores with observers’ scores when scores 2 and 3 were collapsed. The diagonal represents perfect agreement between the true score and observers’ scores. The overall exact agreement with the true scores increased to 78.38% and overscoring reduced to 4.73%.

The average-measure intraclass correlation coefficient (ICC) was 0.91 (95% CI: 0.85 – 0.95, p < 0.0001) among the four observers. The agreement between the individual observers and the true score of WF ranged from moderate to almost perfect. Weighted Cohen’s Kappa and ICC values ranged from 0.62 to 0.82 and 0.66 to 0.82, respectively. The correlation between individual observer score and true score ranged from 0.66 to 0.82 (Table 4). For accuracy, the root mean square error values ranged from 0.55 to 0.87. Mean bias values were generally small (−0.41 – 0.03, Table 4). Observer 1 was the most reliable rater with the highest weighted Cohen’s Kappa, ICC, and correlation coefficient (r), and also the least root mean square error and mean bias.

**Table 4:** Observer agreement and accuracy of scoring window fullness.

| **Observer** | **Kappa** | **RMSE** | **ICC** | **Mean bias** | **r** |
|---|---|---|---|---|---|
| 1 | 0.82 (0.82) | 0.55 (0.33) | 0.82 (0.83) | 0.03 (0.00) | 0.82 (0.83) |
| 2 | 0.65 (0.57) | 0.77 (0.52) | 0.66 (0.58) | -0.16 (-0.05) | 0.66 (0.58) |
| 3 | 0.62 (0.42) | 0.87 (0.68) | 0.66 (0.46) | -0.38 (-0.24) | 0.66 (0.46) |
| 4 | 0.73 (0.63) | 0.72 (0.52) | 0.80 (0.70) | -0.41 (-0.27) | 0.80 (0.70) |

Kappa: weighted Cohen’s Kappa, RMSE: root mean square error, ICC: intraclass correlation coefficient, r: correlation between rater’s score and true score. Values in parentheses are the agreement and accuracy values when scores 2 and 3 were collapsed.

When scores 2 and 3 were collapsed into one score, the average-measure ICC reduced to 0.84 (95% CI: 0.73 – 0.91, p < 0.0001). Collapsing scores 2 and 3 reduced error and bias in scoring (values in parentheses in Table 4). The largest improvement was seen in observers 2 and 3, where their root mean square errors were reduced by 0.25 and 0.19, respectively. Their mean bias also dropped from -0.16 to -0.05 and from -0.38 to -0.24, respectively. However, collapsing scores 2 and 3 reduced the weighted Cohen’s Kappa, ICC, and r of all observers except observer 1 (values in parentheses in Table 4).

### Correlation between WF and absolute and relative lipid content

A positive correlation was observed between WF and absolute lipid content in both sexes (Table 5; male: Spearman correlation = 0.62, p = 0.0079; female: Spearman correlation = 0.58, p = 0.0008). The correlation increased in males (Spearman correlation = 0.69, p = 0.0031), but reduced in females (Spearman correlation = 0.44, p = 0.015), when correlating WF with relative lipid content (Table 5).

**Table 5:** Spearman correlation coefficients between WF and absolute or relative lipid content.

| **Sex** | **Correlation coefficient between WF and absolute lipid content** | **P value of correlation coefficient between WF and absolute lipid content** | **Correlation coefficient between WF and relative lipid mass to wet mass** | **P value of correlation coefficient between WF and relative lipid mass to wet mass** |
|---|---|---|---|---|
| Male | 0.62 | 0.0079 | 0.69 | 0.0031 |
| Female | 0.58 | 0.0008 | 0.44 | 0.0147 |

The fitted regression equations revealed that males with relatively fuller windows had higher absolute and relative lipid contents (Figures 6A and 6B). However, while these variables were still positively correlated in females, fuller windows did not necessarily mean higher absolute or relative lipid content in females (Figures 6A and 6B). However, low WF never incorrectly predicted high absolute or relative lipid content in females.

**Figure 6.**
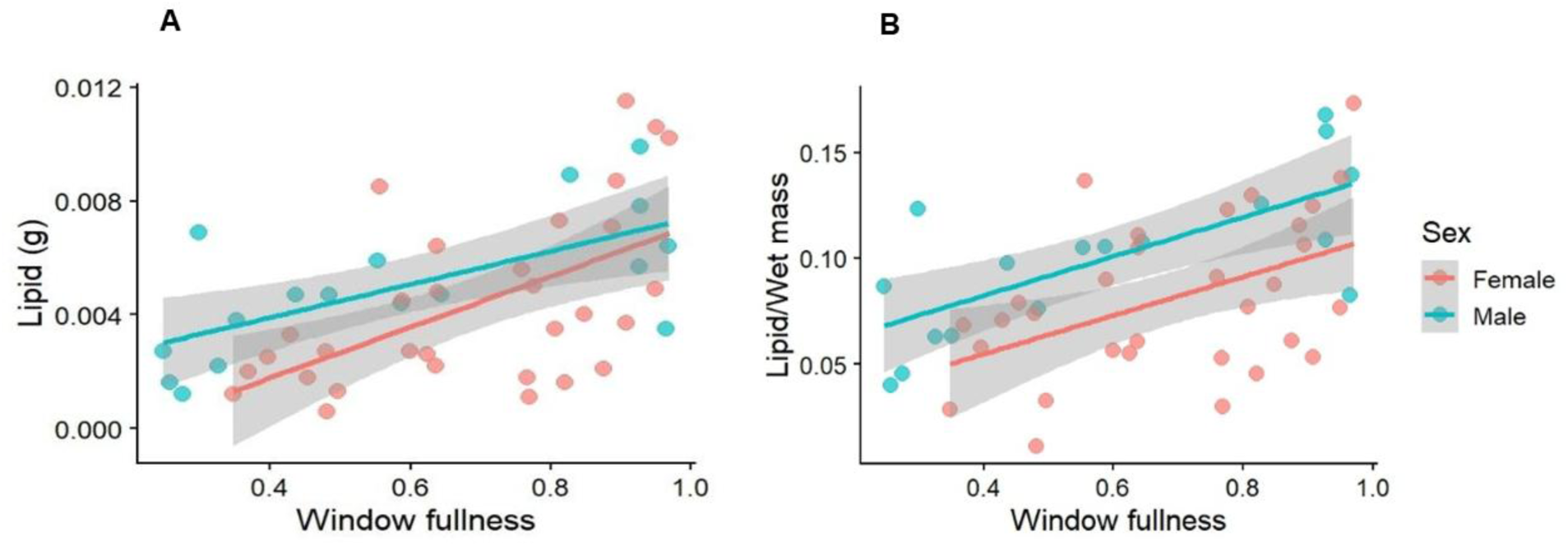
A-B. Relationship between WF and (A) absolute and (B) relative lipid content in male and female BSF. Regression equation for absolute lipid content in males (lipid = 0.0058WF + 0.0016, coefficient of determination = 0.44) and in females (lipid = 0.0089 WF – 0.0018, coefficient of determination = 0.32). Regression equation for relative lipid mass in males (relative lipid mass = 0.093 WF + 0.045, coefficient of determination = 0.51) and in females (relative lipid mass = 0.091 WF + 0.018, coefficient of determination = 0.22).

## Discussion

Pupation is a critical and vulnerable life stage for holometabolous insects, and their eclosion success depends in part on finding an appropriate pupation substrate (Holmes *et al*. 2013). Our data suggest that BSF are able to differentiate between substrates, and prefer the substrate that improves their fitness as adults. Insects have evolved to prefer resources and environments that improve their biological fitness (Gripenberg *et al*. 2010; Ravi Kumar *et al*. 2023) and our data provide further evidence of this phenomenon at the juvenile life stage.

Although pupating in a non-preferred substrate (wood shavings; WS) did not alter survival (eclosion rate) or development time compared to a preferred substrate (potting soil; PS), males and females had smaller dry masses and head widths when pupating in non-preferred substrates, suggesting sublethal impacts of non-preferred pupation medium. Body mass is generally positively correlated with fecundity in adult BSF (Gobbi *et al*. 2013), so smaller body masses can be expected to reduce fitness. Further, females had reduced relative WF (i.e., fat body size), which may reduce resources available for oviposition, as fat reserves are generally used for oviposition by adult female flies (Kalra & Parkash 2014), particularly under energy limitation. Interestingly, thorax length was not affected by pupation substrate, unlike the majority of other morphological measures. The insect thorax is functionally critical for flight in adult flies (Casey *et al*. 2023), including BSF; selection may have kept it stable under both preferred and non-preferred substrates, demonstrating non-isometric body size scaling and resource allocation in the species when exposed to sublethal stressors.

Various pupation substrates are used by BSF producers, including sand, wheat bran, wood shavings, corn cob grits, vermiculite, dry frass, coconut coir dust, potting soil, and sawdust (Holmes *et al*. 2013; Dzepe *et al*. 2020; Boafo *et al*. 2023; Liu *et al*. 2023; Laksanawimol *et al*. 2024). In some cases, pupation substrate may not be provided at all, which can extend development time (Holmes *et al*. 2013). Relatively little research has gone into optimising pupation substrate or determining how pupation substrate may interact with other stressors like low moisture content or handling (but see: Dzepe *et al*. 2020; Durosaro *et al*. 2026). Our study demonstrates a simple method for prepupal pupation substrate preference that could be used to assess a variety of substrate characteristics to maximise fitness and minimize stress.

BSF prepupae in our study strongly preferred PS and sand (SD) to WS, while preferring PS and SD alike. We confirmed that this difference was not related to the looser texture of WS letting in more light, as providing a dark environment did not change the preference for SD over WS. The large particle size of WS may have left prepupae with a sense of ‘exposure’ instead of concealment when compared to finer-grained substrates like PS and SD (Brady & Weil 2008; Vítěz & Trávníček 2010). The rough edges of the WS could also have led to stronger avoidance. Pietrantuono *et al*. (2015) reported avoidance of volcanic ash as a pupation substrate by sawflies (*Notofenusa surosa,* Hymenoptera: Tenthredinidae) because of its sharp edges and abrasive nature.

Generally, the choice of pupation substrates in insects depends on the chemical and physical properties of the substrates, such as porosity, pH, organic matter concentrations, electrical conductivity, and transparency (Prudic *et al*. 2005). Parodi *et al*. (2020) also reported that BSF larvae can differentiate between substrates with different chemicals and microbial contents; presumably, these sensory abilities are retained as prepupae. Physical and chemical properties can be expected to differ significantly in soil- or SD-like substrates and WS. The avoidance of WS as a pupation substrate might also be caused by the presence of some harmful chemicals, such as tannins and resins, which are harmful to insects and can be present in some WS (Fengel & Wegener 2011; Aristri *et al*. 2021). WS might have leached these chemicals (Keržič *et al*. 2024), making BSF prepupae avoid WS as pupation substrate. A future study should examine optimal substrate physical and chemical properties for BSF.

Animals may experience better welfare when they are provided with choices (Dawkins 2021), giving them the liberty to explore and select the preferred option for their health, welfare and/or fitness (Dawkins 2021; Decker *et al*. 2023). A few animals in our study did prefer WS over SD and PS; for these animals, providing a choice of substrate instead of simply recommending that producers provide access to the preferred substrates of the average BSF, could further improve welfare. Further investigation of individual variation in preference in BSF and its relation to health, well-being, and fitness is needed.

Our data also provide moderate support for the use of WF as an indicator of body condition, and thus operational indicator of welfare, in newly-emerged adult BSF. Females had increased WF as adults after pupating in preferred substrates, and a prior study showed that WF declined when females were subjected to juvenile handling stress (Durosaro *et al*. 2026). Lipids stored in the insect fat body are an important energy reserve, sustaining metabolism when food is not available (Ballard *et al*. 2008; Lee & Jang 2014) which is the case for most industry-reared BSF (Barrett *et al*. 2023). However, our study found a strong correlation between WF and total or relative body lipid content only in male BSF, suggesting that, in females, either 1) visible fat mass does not track total fat mass; or 2) fats in other tissues interfered with the correlation.

Importantly, the size of the fat body visible in the ventral abdominal window of adult female BSF may be impacted by the size of other tissues – such as reproductive tissue – located more posteriorly, thus pushing more fat body into the view of the ventral abdominal window and artificially inflating WF score relative to fat body size. This possibility is supported by the WF v. lipid content data we collected: a low WF never *incorrectly predicted* high lipid content – but low lipid content could be observed with a higher WF score. Therefore, low WF may reliably indicate poor lipid content in both females and males and serve as an operational indicator of poor welfare - but high WF does not reliably indicate high relative or absolute lipid content (in females) and better welfare. WF is also impacted by the interaction between sex and age, and sex and mating status (Harjoko *et al*. 2023), highlighting the need to standardize the time and mating status of adult BSF assessed using this measure (as we have done by assessing only 0 – 24 h old, unmated individuals).

WF has potential to be a feasible welfare indicator for use on farms, with good reliability. The rules of thumb for calculating observers’ reliability using at least 30 samples and a minimum of 3 raters proposed by Koo & Li (2016) were followed in this study. An intraclass correlation of 0.5 to 0.75 is indicative of moderate reliability, while 0.75 to 0.9 is indicative of good reliability (Portney & Watkins 2009; Lee *et al*. 2012). Our results suggest that observers can score WF with moderate to almost perfect agreement. Further, the root mean square error and mean bias provide complementary information about the accuracy of the four observers when scoring the WF. The root mean square error and mean bias values obtained for each observer were small, indicating that most observers’ scores were close to the true WF scores, with minimal systematic overestimation or underestimation. The most experienced observer performed best on all metrics, highlighting that training and experience improved reliability, consistency, and accuracy.

We observed that the four observers could not really distinguish between scores 2 and 3 effectively by eye. When we collapsed the indistinguishable scores (scores 2 and 3) into a single category, the mean bias and root mean square error of each of the four observers reduced, indicating improved accuracy, and the average exact agreement increased from 60.14% to 78.38%, indicating that this modified scale provided a clearer and more practical framework for assessing WF in BSF. Our findings on reduced errors when scores 2 and 3 were collapsed support reports by other researchers that reducing the number of ordinal categories can improve reliability (e.g. Stelmer 2004; Meagher 2009).

Despite the improvement in scoring accuracy of WF when scores 2 and 3 were collapsed, a consistent underscoring remained. Such underscoring could have resulted from conservative judgments, and this implied that the observers were more cautious, possibly due to individual or training-based differences in perception thresholds. The dominance of underscoring also highlights the potential role of observer calibration in improving the scoring of BSF WF. Training BSF farmers with standardized image sets or using digital reference scales may improve the accuracy when scoring WF on BSF farms. Although collapsing scores 2 and 3 reduced scoring error by the observers, weighted Cohen’s Kappa, intraclass correlation, and Spearman correlation reduced slightly after collapsing these two scores. This is not surprising because these relative agreement indices are sensitive to score variability (de Raadt *et al*. 2021).

Reducing the number of scores from 5 to 4 reduced the range of scores and diminished the ability of these indices to capture rank-order differences among individuals (Parker *et al*. 2013). Future work may want to continue to finesse the scoring system to see if increased accuracy can be obtained, particularly at the lower end of the WF scale that better predicts low lipid scores in both males and females.

The results from this study and a former study by Durosaro *et al*. (2026) revealed that pupating in non-preferred substrate and handling stress only affected female BSF WF and not male WF on the day of eclosion. The reasons for sex-specific variation in WF shifts associated with stress are not clear - but, a possible explanation is that males may have fewer ‘lipids to burn’, altering their energetic metabolism in response to stress. Alternatively, males that do burn lipids in response to stress may die - ending up in the population of flies whose WF were never assessed in our study. Overall, these results suggest that WF may be an indicator of pre-adult stress in female BSF. A future study should examine how WF affects survivability, behavior, and fecundity.

## Conclusions

BSF prepupae prefer to pupate in soil- or SD-like substrates over WS, and pupation substrate preference has adaptive value, increasing head width and dry mass in flies. From a welfare assessment perspective, WF can be reliably scored as a 4-level ordinal (scores 0 to 3) welfare indicator but further refinement of the scale could make the indicator more precise and valuable. Still, the 4-level WF scale balances feasibility, observer reliability, and construct biological validity, making it promising for operational use in welfare monitoring protocols for adult BSF. WF reliably predicts total and relative fat mass in males, but only successfully predicts low lipid levels in females; therefore, low WF can be used to confidently determine poorer body condition but high WF may not reliably predict better body condition in females. Further research is necessary to link WF to affective state in BSF, similar to other body condition scoring methods used in farmed vertebrate animals.

### Animal welfare implications

The results from our study provide evidence that pupation substrates may have important effects on the welfare of BSF. Prepupae preferred sand and potting soil to wood shavings, and pupating in non-preferred pupation substrates reduced adult fitness traits such as head width, dry mass, and female window fullness. The 78.38% accuracy in scoring window fullness by the observers, the strong relationship between window fullness and lipid content in male flies, and reduction of female window fullness when BSF pupated in non-preferred substrate (and when stressed from our previous study) is an indication that window fullness may be a cheap, practical, non-invasive measure of energetic condition (in males), and welfare in adult BSF. Body condition scoring of adult BSF using window fullness at eclosion time may be suggestive of prior welfare-relevant conditions in BSF.

## Acknowledgement

Authors are grateful to the observers, including Alexander Glica and Rodrigo Rodriguez-Guevara, who scored the window fullness of the flies.

## Author’s contribution

SOD: methodology, investigation, and writing. MB: conceptualization, methodology, investigation, supervision, and writing.

## References

Amrul, N. F., Kabir Ahmad, I., Ahmad Basri, N. E., Suja, F., Abdul Jalil, N. A., & Azman, N. A. (2022). A review of organic waste treatment using black soldier fly (Hermetia illucens). Sustainability, 14(8), 4565. 10.3390/su14084565.

Aristri, M.A., Lubis, M.A.R., Iswanto, A.H., Fatriasari, W., Sari, R.K., Antov, P., Gajtanska, M., Papadopoulos, A.N. & Pizzi, A. (2021). Bio-based polyurethane resins derived from tannin: Source, synthesis, characterisation, and application. Forests, 12(11), 1516. 10.3390/f12111516.

Ballard, J. W. O., Melvin, R. G., & Simpson, S. J. (2008). Starvation resistance is positively correlated with body lipid proportion in five wild caught Drosophila simulans populations. Journal of insect physiology, 54(9), 1371–1376. 10.1016/j.jinsphys.2008.07.009.

Barrett, M., Chia, S.Y., Fischer, B., & Tomberlin, J.K. (2023). Welfare considerations for farming black soldier flies, Hermetia illucens (Diptera: Stratiomyidae): A model for the insects as food and feed industry. Journal of Insects as Food and Feed, 9(2), 119–148. 10.3920/JIFF2022.0041.

Barrett, M., Patel, N., McCarry, B., Shellenberger, G.K., Schwartz, E.R., Fiocca, K., & Waddell, E.A. (2025). Dietary preferences and impacts of feeding on behaviour, longevity, and reproduction in adult black soldier flies (Diptera: Stratiomyidae; Hermetia illucens). Journal of Insects as Food and Feed, 12(4), 691–702. 10.1163/23524588-bja10295.

Boafo, H.A., Gbemavo, D.S.J.C., Timpong-Jones, E.C., Eziah, V., Billah, M., Chia, S.Y., Aidoo, O.F., Clottey, V.A. & Kenis, M., 2023. Substrates most preferred for black soldier fly Hermetia illucens (L.) oviposition are not the most suitable for their larval development. Journal of Insects as Food and Feed, 9(2), 183–192. 10.3920/JIFF2022.0034.

Borkent, S., & Hodge, S. (2021). Glasshouse evaluation of the Black Soldier Fly waste product HexaFrass™ as an organic fertilizer. Insects, 12(11), 977. 10.3390/insects12110977.

Brady, N. C., & Weil, R. R. (2008). The nature and properties of soils (Vol. 13, pp. 662–710). Upper Saddle River, NJ: Prentice Hall.

Casey, C., Heveran, C., & Jankauski, M. (2023). Experimental studies suggest differences in the distribution of thorax elasticity between insects with synchronous and asynchronous musculature. Journal of the Royal Society Interface, 20(201), 20230029. 10.1098/rsif.2023.0029

Dabbou, S., Gai, F., Biasato, I., Capucchio, M.T., Biasibetti, E., Dezzutto, D., Meneguz, M., Plachà, I., Gasco, L., & Schiavone, A. (2018). Black soldier fly defatted meal as a dietary protein source for broiler chickens: Effects on growth performance, blood traits, gut morphology and histological features. Journal of Animal Science and Biotechnology, 9, 1–10. 10.1186/s40104-018-0266-9.

Dawkins, M. S. (2021). The science of animal welfare: Understanding what animals want. Oxford University Press.

de Raadt, A., Warrens, M. J., Bosker, R. J., & Kiers, H. A. (2021). A comparison of reliability coefficients for ordinal rating scales. Journal of Classification, 38(3), 519–543. 10.1007/s00357-021-09386-5.

Decker, S., Lavery, J. M., & Mason, G. J. (2023). Don’t use it? Don’t lose it! Why active use is not required for stimuli, resources or “enrichments” to have welfare value. Zoo Biology, 42(4), 467–475. 10.1002/zoo.21756

Durosaro, S. O., & Barrett, M. (2025). Applying the five domains to farmed black soldier fly, house fly, and yellow mealworm welfare. CABI Reviews, 20(1): 0076. 10.1079/cabireviews.2025.0076.

Durosaro, S.O., Zacarias, M.P., Glica, A., Atkinson, E., Rodriguez-Guevara, R., Jones, E., Gomez, A., & Barrett, M. (2026). Pupation substrate and prepupal handling affect eclosion and adult morphology in black soldier flies. Journal of Insects as Food and Feed (published online ahead of print 2026). 10.1163/23524588-bja10392.

Dzepe, D., Nana, P., Mube, K. H., Fotso, K. A., Tchuinkam, T., & Djouaka, R. (2020). Role of pupation substrate on post-feeding development of black soldier fly larvae, Hermetia illucens (Diptera: Stratiomyidae). Journal of Entomology and Zoology Studies, 8(2), 760–764.

Englund, M. D., & Cronin, K. A. (2023). Choice, control, and animal welfare: definitions and essential inquiries to advance animal welfare science. Frontiers in veterinary science, 10, 1250251. 10.3389/fvets.2023.1250251.

Fengel, D. and Wegener, G. (Eds.). (2011). *Wood: chemistry, ultrastructure, reactions*. Walter de Gruyter. Fisher, M. W. (2020). Pastoral Farming Ethics and Economics–Aligning Grazing Practices and Expectations. Frontiers in Veterinary Science, 7, 511520. 10.3389/fvets.2020.00209.

Fisher, M. W., Schütz, K. E., & Verkerk, G. A. (2020). REVIEW: Critiquing the relationship between body condition and animal welfare. New Zealand Journal of Animal Science and Production, 80, 39–44.

Georgescu, B., Struţi, D., Păpuc, T., Ladoşi, D., & Boaru, A. (2020). Body weight loss of black soldier fly Hermetia illucens (Diptera: Stratiomyidae) during development in non-feeding stages: Implications for egg clutch parameters. European Journal of Entomology, 117, 216–225. 10.14411/eje.2020.023.

Ghosh, C. P., Datta, S., Mandal, D., Das, A. K., Roy, D. C., Roy, A., & Tudu, N. K. (2019). Body condition scoring in goat: Impact and significance. Journal of Entomology and Zoology Studies, 7(2), 554–560.

Gobbi, P., Martínez-Sánchez, A. & Rojo, S. (2013) The effects of larval diet on adult life-history traits of the black soldier fly, *Hermetia illucens* (Diptera: Stratiomyidae). European Journal of Entomology, 110, 461–468. 10.14411/eje.2013.061.

Gripenberg, S., Mayhew, P. J., Parnell, M., & Roslin, T. (2010). A meta-analysis of preference– performance relationships in phytophagous insects. Ecology letters, 13(3), 383–393. 10.1111/j.1461-0248.2009.01433.x.

Harjoko, D. N., Hua, Q. Q. H., Toh, E. M. C., Goh, C. Y. J., & Puniamoorthy, N. (2023). A window into fly sex: Mating increases female but reduces male longevity in black soldier flies. Animal Behaviour, 200, 25–36. 10.1016/j.anbehav.2023.03.007.

Hogsette J. A. (1992). New diets for production of house flies and stable flies (Diptera: Muscidae) in the laboratory. Journal of economic entomology, 85(6), 2291–2294. 10.1093/jee/85.6.2291.

Holmes, L. A., Vanlaerhoven, S. L., & Tomberlin, J. K. (2013). Substrate effects on pupation and adult emergence of Hermetia illucens (Diptera: Stratiomyidae). Environmental entomology, 42(2), 370–374. 10.1603/EN12255.

Johnson, M. & Barrett, M. (2025). Review: Exploring correctness, usefulness, and feasibility of potential physiological operational welfare indicators for farmed insects to establish research priorities. Animal, 101501. 10.1016/j.animal.2025.101501.

Kalra, B., & Parkash, R. (2014). Trade-off of ovarian lipids and total body lipids for fecundity and starvation resistance in tropical populations of Drosophila melanogaster. Journal of Evolutionary Biology, 27(11), 2371–2385. 10.1111/jeb.12480.

Kaya C, Generalovic TN, Ståhls G, Hauser M, Samayoa AC, Nunes-Silva CG, Roxburgh H, Wohlfahrt J, Ewusie EA, Kenis M, Hanboonsong Y, Orozco J, Carrejo N, Nakamura S, Gasco L, Rojo S, Tanga CM, Meier R, Rhode C, Picard CJ, Jiggins CD, Leiber F, Tomberlin JK, Hasselmann M, Blanckenhorn WU, Kapun M, Sandrock C. (2021). Global population genetic structure and demographic trajectories of the black soldier fly, Hermetia illucens. BMC Biology. 19(1): 94. 10.1186/s12915-021-01029-w.

Keržič, E., Vek, V., Oven, P., & Humar, M. (2024). Changes in wood durability due to leaching of biologically active substances (extractives) resulting from weathering. Case Studies in Construction Materials, 21, e03921. 10.1016/j.cscm.2024.e03921.

Kim, Y. B., Kim, D. H., Jeong, S. B., Lee, J. W., Kim, T. H., Lee, H. G., & Lee, K. W. (2020). Black soldier fly larvae oil as an alternative fat source in broiler nutrition. Poultry Science, 99(6), 3133–3143. 10.1016/j.psj.2020.01.018.

Koo, T. K., & Li, M. Y. (2016). A guideline of selecting and reporting intraclass correlation coefficients for reliability research. Journal of chiropractic medicine, 15(2), 155–163. 10.1016/j.jcm.2016.02.012.

Kortsmit, Y., Van der Bruggen, M., Wertheim, B., Dicke, M., Beukeboom, L. W., & Van Loon, J. J. A. (2023). Behaviour of two fly species reared for livestock feed: optimising production and insect welfare. Journal of Insects as Food and Feed, 9(2), 149–169. 10.3920/JIFF2021.0214.

Laksanawimol, P., Anukun, P., & Thancharoen, A. (2024). Use of different dry materials to control the moisture in a black soldier fly (Hermetia illucens) rearing substrate. PeerJ, 12, e17129. 10.7717/peerj.17129.

Lee, K. P., & Jang, T. (2014). Exploring the nutritional basis of starvation resistance in Drosophila melanogaster. Functional Ecology, 28(5), 1144–1155. 10.1111/1365-2435.12247.

Lee, K.M., Lee, J., Chung, C.Y., Ahn, S., Sung, K.H., Kim, T.W., Lee, H.J., & Park, M.S. (2012). Pitfalls and important issues in testing reliability using intraclass correlation coefficients in orthopaedic research. Clinics in orthopedic surgery, 4(2), 149–155. 10.4055/cios.2012.4.2.149.

Leong, S. Y., Kutty, S. R. M., Malakahmad, A., & Tan, C. K. (2016). Feasibility study of biodiesel production using lipids of Hermetia illucens larva fed with organic waste. Waste Management, 47, 84–90. 10.1016/j.wasman.2015.03.030.

Lindholm-Perry, A. K., Kuehn, L. A., Freetly, H. C., Oliver, W. T., Neville, B. W., Crouse, M. S., Thorson, J. F., Wells, J. E., & Keel, B. N. (2024). 262 Why are some cows skinny? An evaluation of stress and immune factors. Journal of Animal Science, 102(Suppl 3), 88. 10.1093/jas/skae234.099.

Liu, Z., Morel, P. C. H., & Minor, M. A. (2023). Substrate and moisture content effects on pupation of the black soldier fly (Diptera: Stratiomyidae). Journal of Insects as Food and Feed, 9(4), 415–425. 10.3920/JIFF2022.0057.

Marshall, S. A., Woodley, N. E., & Hauser, M. (2015). The historical spread of the black soldier fly, Hermetia illucens (L.) (Diptera, Stratiomyidae, Hermetiinae), and its establishment in Canada. The Journal of the Entomological Society of Ontario, 146. 10.21083/jeso.v146i0.3696.

Matthews, L., Cameron, C., Sheahan, A., Kolver, E., & Roche, J. (2012). Associations among dairy cow body condition and welfare-associated behavioral traits. Journal of Dairy Science, 95(5), 2595–2601. 10.3168/jds.2011-4889.

McKay, H. and Shah, S., 2025. Forecasting farmed animal numbers in 2033. Rethink Priorities. Available at https://rethinkpriorities.org/research-area/forecasting-farmed-animal-numbers-in-2033/.

Meagher, R. K. (2009). Observer ratings: Validity and value as a tool for animal welfare research. Applied Animal Behaviour Science, 119(1-2), 1–14. 10.1016/j.applanim.2009.02.026

Nayak, A., Rühl, M., & Klüber, P. (2024). *Hermetia illucens* (Diptera: Stratiomyidae): Need, Potentiality, and Performance Measures. Agriculture, 14(1), 8. 10.3390/agriculture14010008.

Nielsen, S. S., Alvarez, J., Boklund, A., Dippel, S., Dorea, F., Figuerola, J., Herskin, M. S., Michel, V., Miranda Chueca, M. A., Nannoni, E., Nonno, R., Riber, A. B., Stahl, K., Stegeman, J. A., Thulke, H. H., Tuyttens, F., Cozzi, G., Knierim, U., Martí, S., . . . Winckler, C. (2025). Welfare of beef cattle. EFSA Journal, 23(7), e9518. 10.2903/j.efsa.2025.9518

Nogales-Mérida, S., Gobbi, P., Józefiak, D., Mazurkiewicz, J., Dudek, K., Rawski, M., Kierończyk, B. and Józefiak, A. (2019). Insect meals in fish nutrition. Reviews in Aquaculture, 11(4), 1080–1103. 10.1111/raq.12281.

Oliveira, F. R., Doelle, K., & Smith, R. P. (2016). External morphology of Hermetia illucens Stratiomyidae: Diptera (L. 1758) based on electron microscopy. Annual research & review in biology, 9(5), 1–10. 10.9734/ARRB/2016/22973.

Parker, R. I., Vannest, K. J., & Davis, J. L. (2013). Reliability of multi-category rating scales. Journal of School Psychology, 51(2), 217–229. 10.1016/j.jsp.2012.12.003.

Parodi, A., Van Dijk, K., Van Loon, J. J., De Boer, I. J., Van Schelt, J., & Van Zanten, H. H. (2020). Black soldier fly larvae show a stronger preference for manure than for a mass-rearing diet. Journal of Applied Entomology, 144(7), 560–565. 10.1111/jen.12768.

Pietrantuono, A.L., Enriquez, A.S., Fernández-Arhex, V. and Bruzzone, O.A., 2015. Substrates preference for pupation on sawfly Notofenusa surosa (Hymenoptera: Tenthredinidae). Journal of Insect Behavior, 28(3): 257–267. 10.1007/s10905-015-9499-z.

Portney, L. G., & Watkins, M. P. (2009). Foundations of clinical research: applications to practice (Vol. 892, pp. 11–15). Upper Saddle River, NJ: Pearson/Prentice Hall.

Prudic, K. L., Oliver, J. C., & Bowers, M. D. (2005). Soil nutrient effects on oviposition preference, larval performance, and chemical defense of a specialist insect herbivore. Oecologia, 143(4), 578–587. 10.1007/s00442-005-0008-5.

Ravi Kumar, V., Agavekar, G., & Agashei, D. (2023). Fitness landscapes reveal context-dependent benefits of oviposition behavior. Evolution, 77(2), 550–561. 10.1093/evolut/qpac035.

Renggaman, A., Choi, H. L., Sudiarto, S. I., Alasaarela, L., & Nam, O. S. (2015). Development of pig welfare assessment protocol integrating animal-, environment-, and management-based measures. Journal of Animal Science and Technology, 57, 1. 10.1186/s40781-014-0034-0.

Rivers, D.B., Thompson, C. and Brogan, R. (2011). Physiological trade-offs of forming maggot masses by necrophagous flies on vertebrate carrion. Bulletin of Entomological Research 101: 599–611. 10.1017/S0007485311000241.

Roche JR, Friggens NC, Kay JK, Fisher MW, Stafford KJ, Berry DP. (2009). Invited review: Body condition score and its association with dairy cow productivity, health, and welfare. Journal of Dairy Science. 92(12):5769–801. 10.3168/jds.2009-2431.

Schneider, C. A., Rasband, W. S., & Eliceiri, K. W. (2012). NIH Image to ImageJ: 25 years of image analysis. Nature Methods, 9(7), 671–675. 10.1038/nmeth.2089.

Stemler, S. E. (2004). A comparison of consensus, consistency, and measurement approaches to estimating interrater reliability. Practical Assessment, Research, and Evaluation, 9(1).

Tomberlin, J. K. & Van Huis, A. (2020). Black soldier fly from pest to ‘crown jewel’ of the insects as feed industry: an historical perspective. Journal of Insects as Food and Feed, 6(1), 1–4. 10.3920/JIFF2020.0003.

Tomberlin, J. K., Sheppard, D. C., & Joyce, J. A. (2002). Selected life-history traits of black soldier flies (Diptera: Stratiomyidae) reared on three artificial diets. Annals of the Entomological Society of America, 95(3), 379–386. 10.1603/0013-8746(2002)095[0379:SLHTOB]2.0.CO;2.

Vall, E., Blanchard, M., Sib, O., Cormary, B., & González-García, E. (2025). Standardized body condition scoring system for tropical farm animals (large ruminants, small ruminants, and equines). Tropical Animal Health and Production, 57(2), 106. 10.1007/s11250-025-04328-4

Veissier, I., Aubert, A., & Boissy, A. (2012). Animal welfare: A result of animal background and perception of its environment. Animal Frontiers, 2(3), 7–15. 10.2527/af.2012-0043.

Vieira, A., Brandão, S., Monteiro, A., Ajuda, I., & Stilwell, G. (2015). Development and validation of a visual body condition scoring system for dairy goats with picture-based training. Journal of dairy science, 98(9), 6597–6608. 10.3168/jds.2015-9428

Vítěz, T., & Trávníček, P. (2010). Particle size distribution of sawdust and wood shavings mixtures. Research in Agricultural Engineering, 56(4), 154–158. 10.17221/8/2010-RAE

Welfare Quality® (2009). Welfare Quality® assessment protocol for cattle. Welfare Quality® Consortium, Lelystad, Netherlands.

Williams, C. M., Thomas, R. H., MacMillan, H. A., Marshall, K. E., & Sinclair, B. J. (2011). Triacylglyceride measurement in small quantities of homogenised insect tissue: Comparisons and caveats. Journal of Insect Physiology, 57(12), 1602–1613. 10.1016/j.jinsphys.2011.08.008.

Zufferey, R., Minnig, A., Thomann, B., Zwygart, S., Keil, N., Schüpbach, G., Miserez, R., Zanolari, P., & Stucki, D. (2021). Animal-Based Indicators for On-Farm Welfare Assessment in Sheep. Animals : An Open Access Journal From MDPI, 11(10), 2973. 10.3390/ani11102973

