## Supplementary File 1 for "Evaluation of window fullness at time of eclosion as an operational welfare indicator in farmed black soldier flies (*Hermetia illucens*)"

### BSF Window Fullness Training

<u>What is the 'window'?</u>

Black soldier fly adults have a clear window on the ventral (underside) surface of their abdomen.

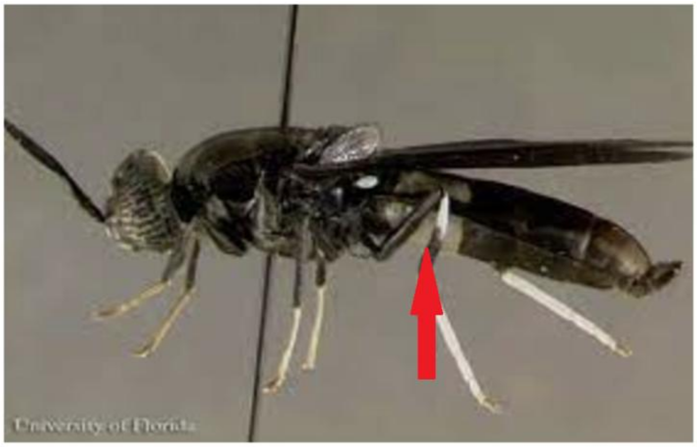


You can see this window from the side view of this male BSF, as indicated by the red arrow. It extends through the first two abdominal terga in both males and females, though there is sometimes variation in the exact patterning of the cuticle's melanisation.

<u>What is the goal of this training?</u>

The goal of this training is to assist you in developing the skills necessary to evaluate when a live BSF adult has a window that is *more* or *less* full as a possible operational welfare indicator to be used on farms. We hope that, by accessing this simple training, you are better prepared to determine when a window is full to a particular percentage. Then, by testing multiple experimenters on live specimens, this will allow us to assess accuracy of experimenters with different levels of familiarity with window fullness analysis and interindividual variation among observers.

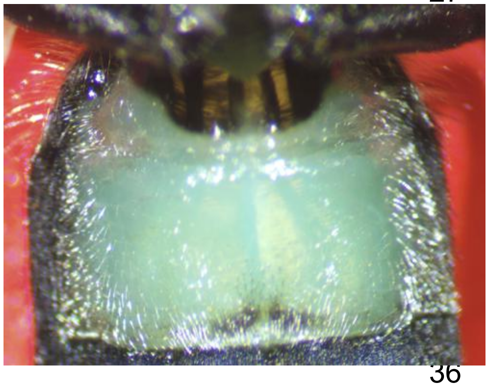

<u>What is the window full of?</u>

This window allows the animal's fat body to be (partially) observed through the clear panel of their cuticle (see below). This generally appears as an opaque, creamy blue-ish to white-ish mass (see photo to the left). We want to know how **full the window is of fat from the fat body.**

Importantly, just as the fat body can be seen through the window, so can other internal anatomical structures such as nerves, trachea (tubes for breathing), or elements of the digestive system. **These do not count as the fat body.** When we are considering how 'full' the window is, we want to consider **only** the fat body and not the digestive system or other anatomical parts.

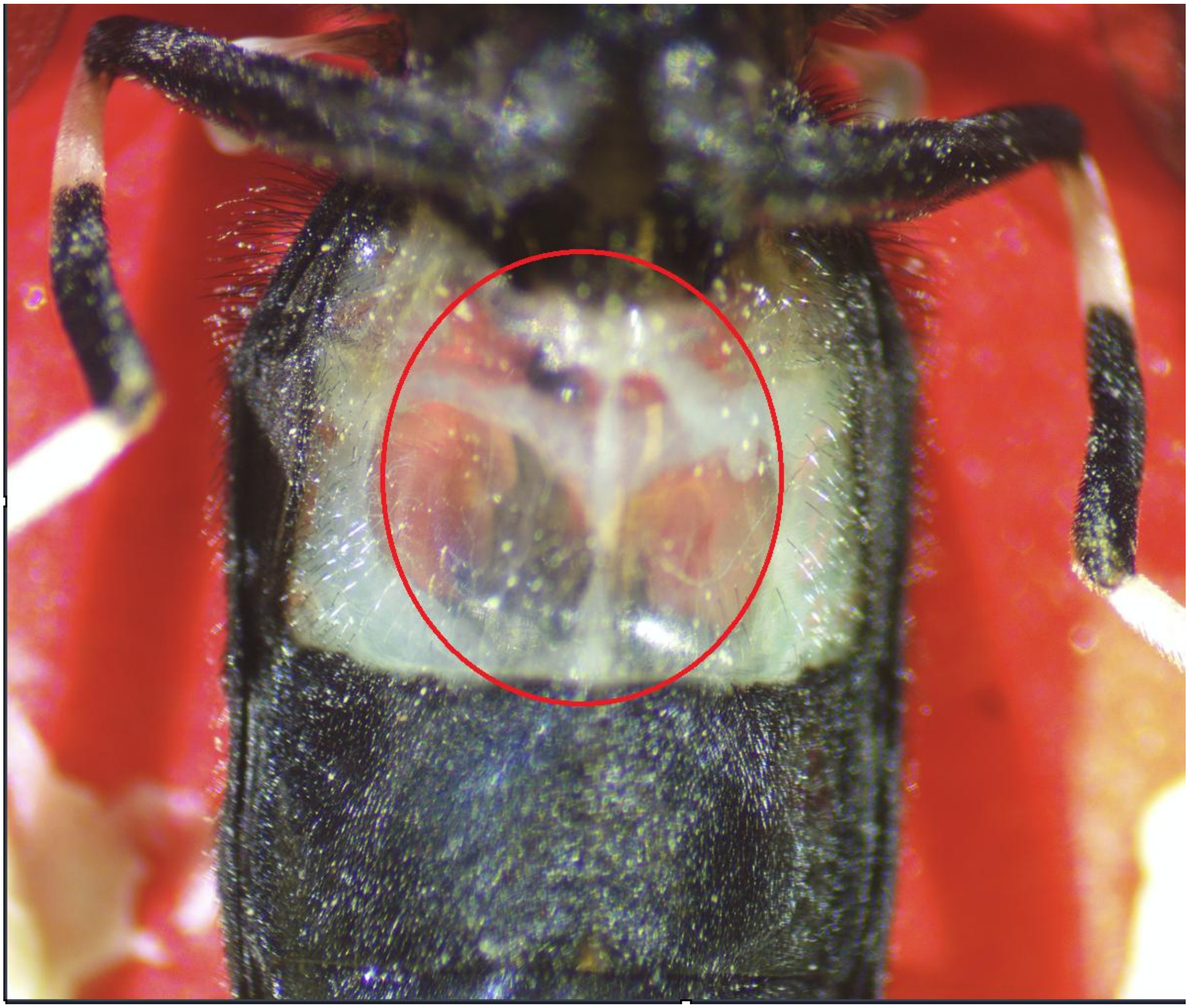

For example, in the above photo, you can see the fat body, which is a bit slim and wispy on either side of the red circle on the exterior margins of the window itself. Within the circle is the digestive system - it is a very wispy, semi-clear structure that has a Y shape. There is a central region that extends vertically down the midline of the body. Then, there are two ‘wings’ that extend to the left and right, around ⅔ of the way up the structure. Finally, at the top is a bit of a triangle that attaches to this element. None of this material is considered fat body and thus doesn’t count in analysing window fullness.

Here is another close-up example; the bottom of the digestive system has now been obscured by fat body, but the middle is still clearly visible as digestive tissue inside the red circle.

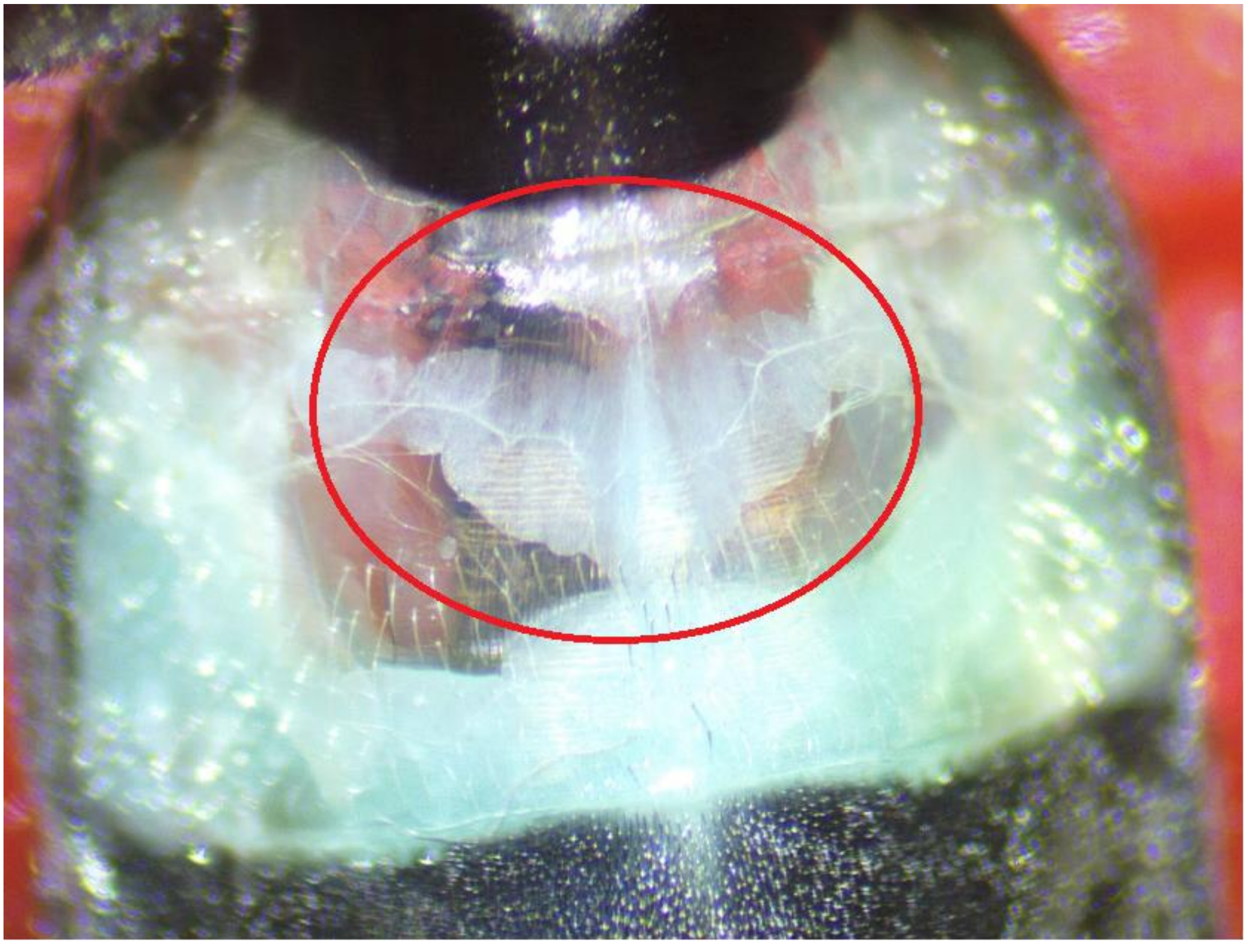

**Windows can be categorized as a ‘0’, ‘1’, ‘2’, ‘3’, or ‘4’.**

- Level 0 = 0-19.99% full of the fat body
- Level 1 = 20%-39.99% full of the fat body
- Level 2 = 40%-59.99% full of the fat body
- Level 3 = 60%-79.99% full of the fat body
- Level 4 = 80%-100% full of the fat body

**Level 0 is extremely uncommon. Here are some examples:**

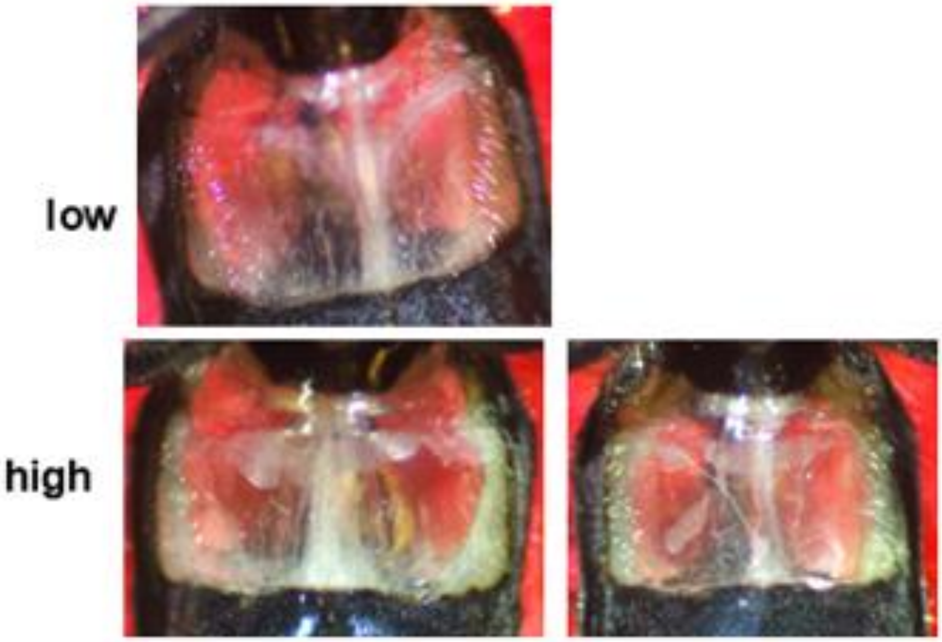


**Level 1 is uncommon. Here are some examples:**

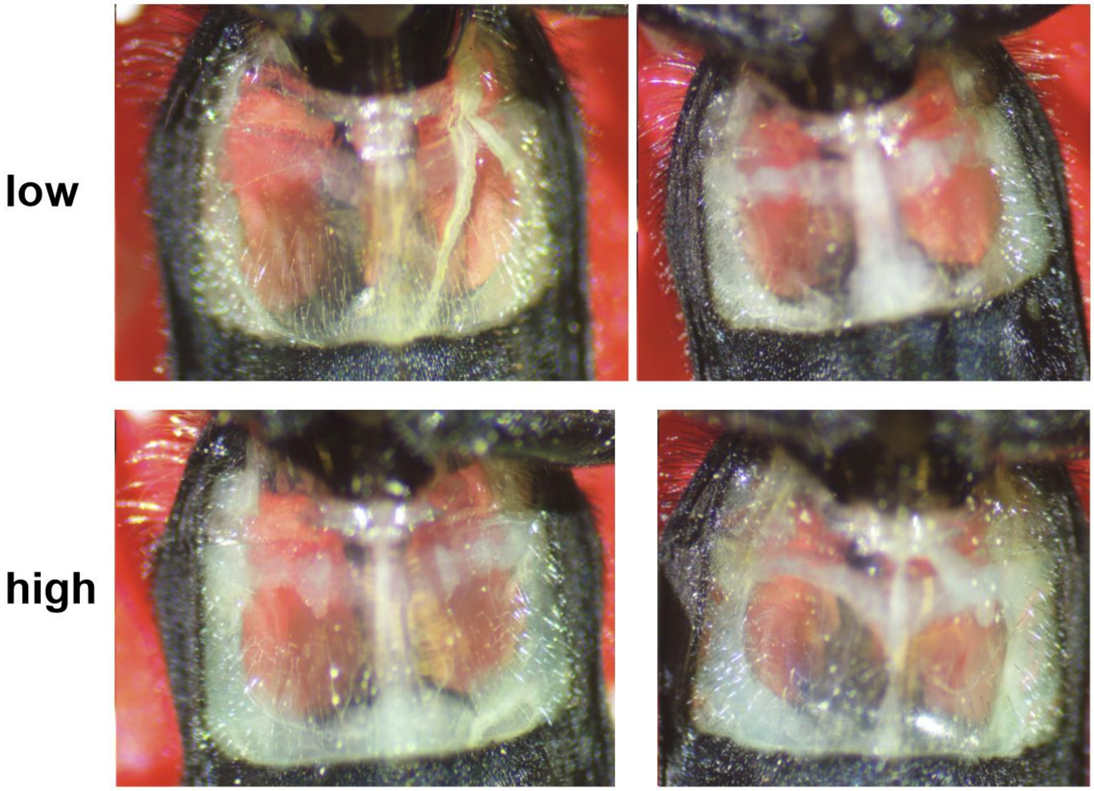

**Level 2 is more common. Here are some examples:**

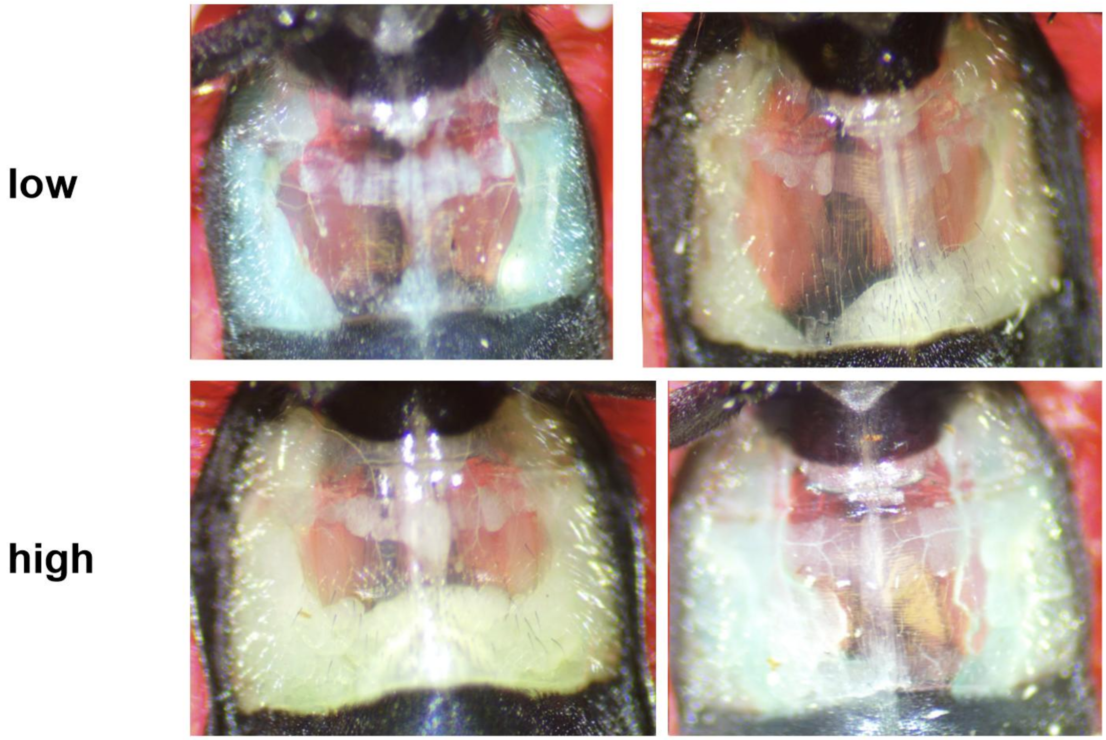


**Level 3 is common. Here are some examples:**

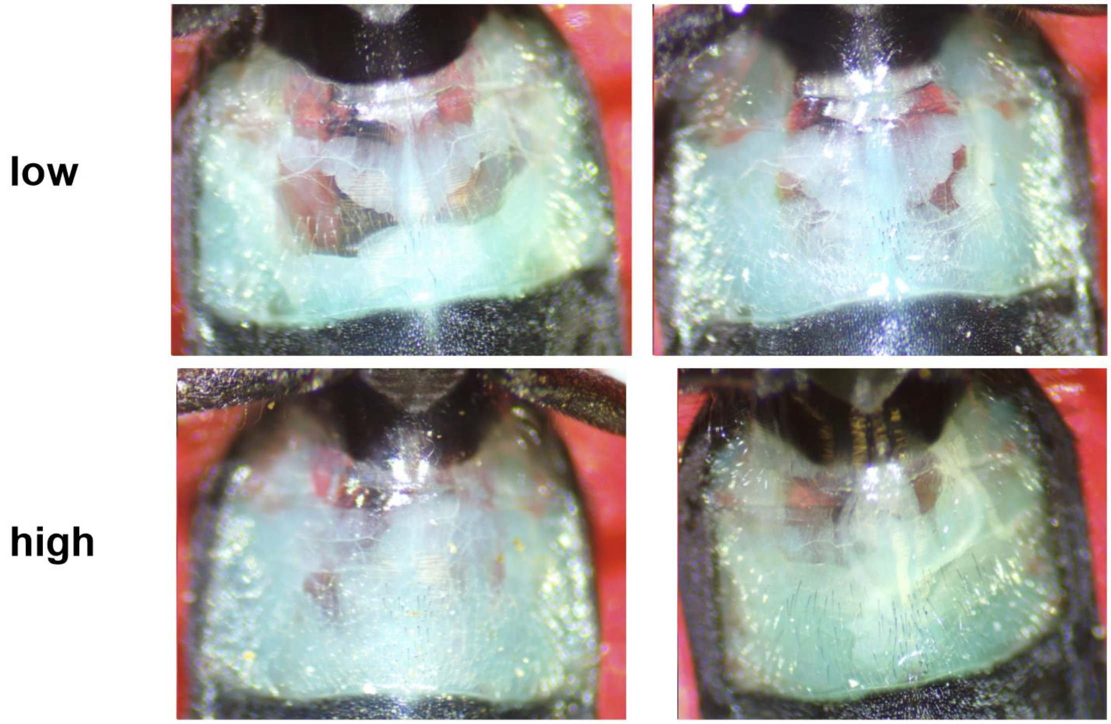

**Level 4 is common. Here are some examples:**

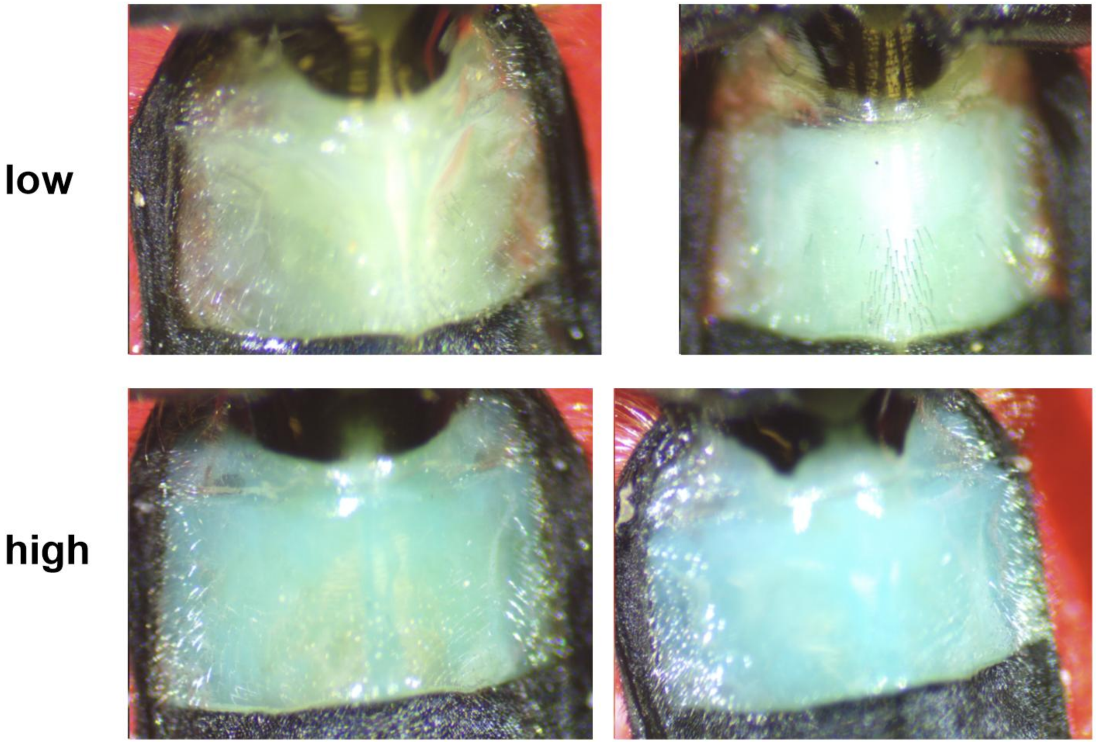
